# PCDH7 Promotes Cell Migration by Regulating Myosin Activity

**DOI:** 10.1101/2021.09.21.460794

**Authors:** Mohammad Haroon Qureshi, Halil Bayraktar, M. Talha Cinko, Cansu Akkaya, Altug Kamacioglu, Z. Cansu Uretmen-Kagiali, Erdem Bozluolcay, Nurhan Ozlu

**Affiliations:** Department of Molecular Biology and Genetics, Koç University, 34450, Istanbul, Turkey; Biomedical Sciences and Engineering Graduate Program, Koç University, 34450, Istanbul, Turkey; Molecular Biology and Genetics Department, Istanbul Technical University, 34469, Istanbul, Turkey; Koç University Research Center for Translational Medicine (KUTTAM), 34450, Istanbul, Turkey

**Keywords:** cell cortex, myosin, protocadherin, cell migration, ERM

## Abstract

Cell migration requires spatiotemporally coordinated activities of multicomponent structures including the actomyosin cortex, plasma membrane, adhesion complexes and the polarity proteins. How they function together to drive this complex dynamic process remains an outstanding question. Here, we show that a member of the protocadherin family, PCDH7 displays a polarized localization in migratory cells with a dynamic enrichment at the leading and rear edges. Perturbation of PCDH7 interferes with the migration of nontransformed retinal pigment epithelial cells and the invasion of cancer cells. The overexpression of PCDH7 enhances the migration capability of cortical neurons *in vivo*. PCDH7 interacts with the myosin phosphatase subunits MYPT1 and PP1cβ. Ectopic expression of PCDH7 enhances the MYPT1 inhibitory phosphorylation levels and the phosphorylation of the myosin regulatory light chain and ERM at the polarized cortex. The chemical inhibition of phosphatase activity recovers migration phenotypes of PCDH7 knockout cells. We propose that PCDH7 regulates phosphorylation thus the activity of myosin and ERM at the polarized cortex through its interaction with myosin phosphatase. Collectively, our study suggests a new component for the spatial coordination of the plasma membrane and the cortex during cell migration.

## Introduction

Cell motility is an indispensable feature of a wide range of biological processes including mammalian embryonic development, inflammatory response, and tissue repair and regeneration (Ridley et al., 2003; Te Boekhorst et al., 2016). Cell motility also features in the invasion and metastasis of tumor cells (Hanahan and Weinberg, 2011; Schwartz and Horwitz, 2006; Svitkina, 2018; Te Boekhorst et al., 2016). During cell migration, actin-based structures, lamellipodia and filopodia form at the leading edge of migrating cells (Cramer, 1999; Cramer, 2013; Cramer et al., 1997). Protrusive forces are generated by those structures through polymerization of actin filaments (Svitkina, 2018) which is regulated by ERM (Ezrin-Radixin-Moesin) proteins (Arpin et al., 2011). Phosphorylation of ERM by various kinases stimulates lamellipodium and filopodium formation thus membrane protrusion and cortical stability through its simultaneous interaction with F-actin and plasma membrane (Baumgartner et al., 2006; Hipfner et al., 2004; Ponti et al., 2004; Uretmen Kagiali et al., 2020).

The contractility of the actomyosin cytoskeleton is mediated by the motor activity of myosins, mainly myosin II, also known as the non-muscle myosin II (NMII), which plays a central role in cell migration (Vicente-Manzanares et al., 2009). The Myosin II-based force generation is dynamically controlled through the phosphorylation of its light chain, Myosin Light Chain 2 (MLC2)/ Regulatory Light Chain (MRLC). Phosphorylation of MLC2 at Serine19 (S19) reversibly drives the ATPase activity of myosin II by enhancing the Mg^2+^-ATPase activity of the complex in the presence of actin (Houdusse and Sweeney, 2016; Somlyo and Somlyo, 2003). Phosphorylation of MLC2 at Threonine18 (T18) further enhances the Mg^2+^-ATPase activity of myosin II (Ikebe et al., 1986). Phosphorylation of MLC2 triggers myosin’s binding to actin and can be governed by the activity of more than a dozen kinases including MLCK, ROCK, ZIPK, CDC42BP (Vicente-Manzanares et al., 2009). Similarly, MLC2 dephosphorylation by phosphatases such as myosin phosphatase also regulates the motor activity of the myosin complex (Ito et al., 2004; Kiss et al., 2019). Myosin motors-based contractility plays a central role in all steps of cell migration through its interaction with the actin cytoskeleton. Although its impact in retraction, protrusion and adhesion of a migratory cell is well known (Cai et al., 2006; Juanes-Garcia et al., 2015; Schwartz and Horwitz, 2006; Vicente-Manzanares et al., 2008; Vicente-Manzanares et al., 2009; Vicente-Manzanares et al., 2007), the regulation of the complex interplay between myosin and other cortical and plasma membrane elements is incompletely understood.

Members of the cadherin superfamily exhibit differential roles in the development and progression of cancer (Jeanes et al., 2008; Mendonsa et al., 2018; Mrozik et al., 2018). Members of the largest subfamily of the cadherin superfamily, namely protocadherins, are widely expressed in the nervous system. Their functions are poorly explored but they are gradually being shown to play important roles in cell-cell interactions in the nervous system (Hayashi and Takeichi, 2015). Non-clustered protocadherins, a subset of protocadherins, have recently been suggested to play important roles in neuronal functions and cell migration. Unlike the classical cadherins, their cell adhesion role is context-dependent, as they can also exhibit anti-adhesion roles during development and cell division (Jontes, 2016; Ozlu et al., 2015; Qiu et al., 2016; Yasuda et al., 2007; Zhang et al., 2012; Zhou et al., 2017; Zhu et al., 2014). One such member, PCDH7, belongs to the δ1 subclass of non-clustered protocadherins. Its roles in mammalian cells are not well characterized. PCDH7 has been reported to be highly expressed in brain metastasizing breast cancer cell lines (Bos et al., 2009). PCDH7 mediated metastasis of breast cancer cells to the brain are primarily driven by cell-cell contacts between astrocytes and invading cancer cells. PCDH7 stabilizes connexin-43 mediated gap junctions and drives STING signaling across the cells, thus enhances the survival of invading cells (Chen et al., 2016). A Xenopus homolog of PCDH7, NF-protocadherin (NF-PC) has been previously shown to play a role in ectoderm differentiation and retinal axon growth and guidance through its interaction with TAF1/SET (Heggem and Bradley, 2003; Leung et al., 2015; Leung et al., 2013; Piper et al., 2008). TAF1/SET is an inhibitor of phosphatase, PP2A. PCDH7 has four different isoforms, which have identical extracellular and transmembrane domains but differ in their cytosolic domains (**Figure 1A**). PCDH7a and PCDH7b have smaller cytosolic domains, whereas PCDH7c and PCDH7d have longer cytosolic domains with an additional PP1α interaction motif (RRVTF) in its cytoplasmic domain (CM3) (Yoshida et al., 1999). PCDH7 has been implicated in lung tumorigenesis by inducing MAPK signaling through SET mediated PP2A inhibition (Zhou et al., 2017). All isoforms share SET binding domain and overexpression of all four isoforms can transform human bronchial epithelial cells (Zhou et al., 2017).

**Figure 1.**
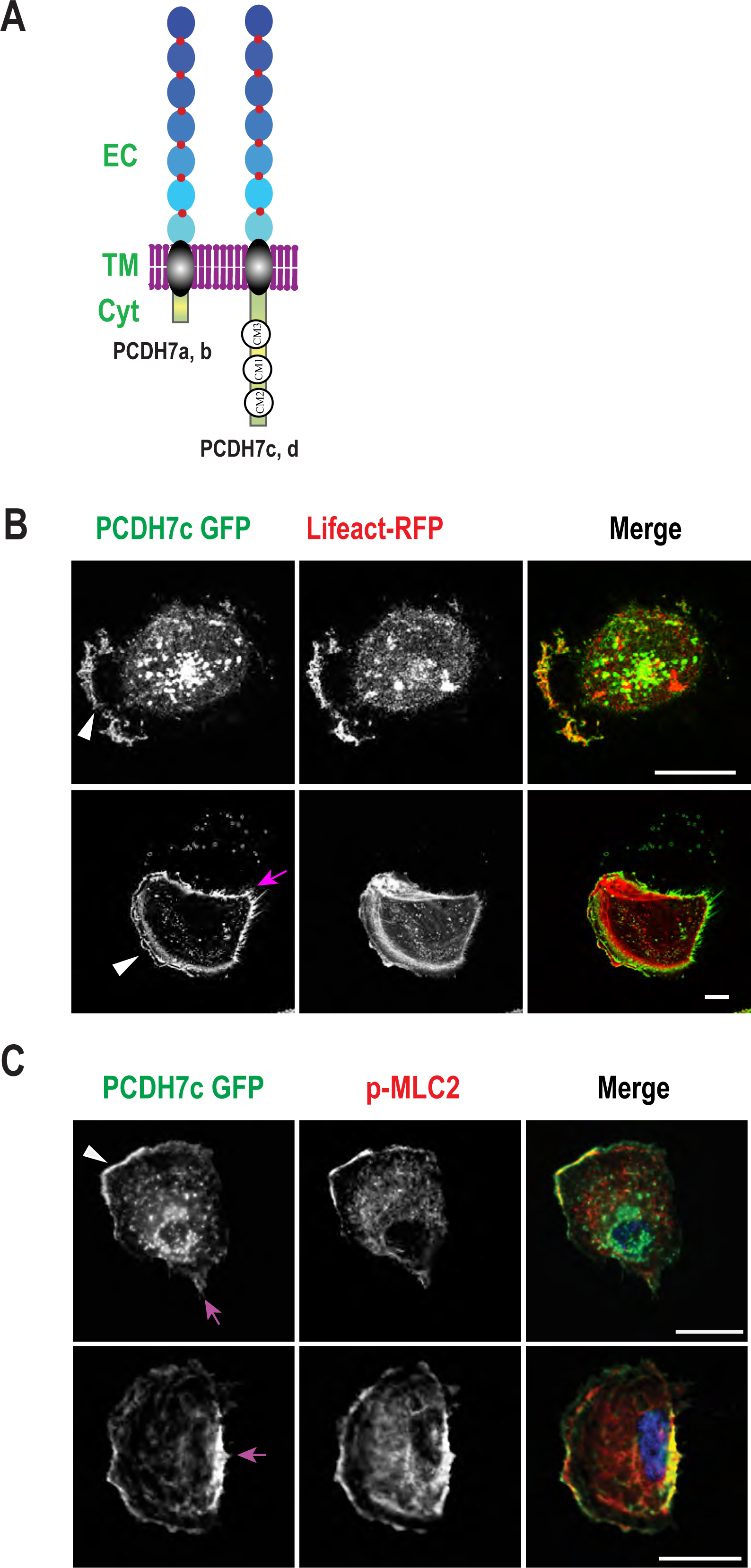
PCDH7 localizes at the protruding front and trailing rear edge of migratory RPE1 cells. **(A)** A diagram showing PCDH7 isoforms and their domains: Extracellular domain (EC), transmembrane domain (TM), and cytosolic domain (Cyt) including CM1, CM2 and CM3 domains in longer isoforms (PCDH7c and d). (B) Representative images from live imaging of RPE1 cells expressing PCDH7-GFP (green) and lifeact-RFP (red). Upper panel shows a migrating RPE1 cell expressing PCDH7 GFP (green) and lifeact RFP (red) with PCDH7 enrichment on protruding front edge (white arrow) (video 1). Lower panel shows a still image from video 2 with PCDH7 enrichment on the cell rear (magenta arrow). **(C)** Immunostaining of RPE1 cells expressing PCDH7-GFP are visualized using anti-GFP (green), anti-phosho-Myosin Light Chain 2 (p-MLC2, S19) (red) antibodies. Colocalization of PCDH7 and p-MLC2 at the front (white arrow, top) and rear (magenta arrow, bottom) edges. Scale bars: 20 μm.

Our previous work has shown that PCDH7 enriches and localizes all over the cell surface during mitosis, but it is largely restricted to cell-cell contact regions in interphase cells (Ozlu et al., 2015). Perturbation of PCDH7 expression affected cell rounding pressure of mitotic cells, an effect was also seen upon myosin IIA heavy chain (MYH9) perturbation (Ozlu et al., 2015). Despite the emerging role of PCDH7 in the cellular cortex, its molecular mechanism in mammalian cells remains to be uncovered. In this study, we explore molecular details of the function of PCDH7 in cell migration. Our work shows that PCDH7 is associated with cortical actomyosin with a dynamic enrichment at the leading and rear edges. RPE1 cells lose their persistence of migration upon PCDH7 perturbation, an effect that is recovered upon PCDH7 replenishment. We showed that PCDH7 interacts with the myosin phosphatase subunits PP1cβ and MYPT1. PCDH7 expression is correlated with phosphorylation of the myosin regulatory light chain and ERM at the polarized cortex. We propose that PCDH7 promotes cell migration by enabling phosphorylated myosin complexes and ERM to establish mechanical properties of the polarized cortex during migration.

## Results

### PCDH7 associates with cortical actomyosin and localizes to leading and rear edges in migrating RPE1 cells

As a member of the cadherin superfamily, many protocadherins are known to be connected to the cortical actomyosin cytoskeleton (Budnar and Yap, 2013; Hayashi et al., 2014; Moeller et al., 2004; Ramakrishnan et al., 2012; Ratheesh and Yap, 2012). To investigate the association of PCDH7 with the actomyosin cortex during migration, we used Retinal Pigment Epithelial cells (RPE1) which are non-transformed migratory human cells (Jin et al., 2000). Isoform C of PCDH7-GFP, (PCDH7c-GFP) and lifeact-RFP fusion proteins expressing RPE1 cells were imaged using time-lapse confocal scanning microscopy. In a migrating RPE1 cell, PCDH7c-GFP is enriched on the protruding leading edge as the cell displayed random migration **(Video 1,** **Figure 1B, top panel)**. In addition to the enrichment at the leading edge, PCDH7c also co-localizes with F-actin **(Video 2,** **Figure 1B, bottom panel)** at the cell rear edge. PCDH7 displays a dominant vesicular presence inside the cell, owing to its enrichment in the endoplasmic reticulum as shown by our previous study (Ozlu et al., 2015). The shorter isoform of PCDH7 (PCDH7b) also displayed similar subcellular localization. Albeit PCDH7b showed a higher predominance in the vesicles compared to that of the longer isoform (PCDH7c) **(Video 3)**.

During actin-based cell migration, myosin II enriches at the rear edge and moves the cell forward by its motor activity causing rear edge retraction (Conti and Adelstein, 2008; Cramer, 2013; Vicente-Manzanares et al., 2009). Next, we examined the localization of PCDH7 and active myosin by imaging PCDH7 GFP and pMLC2 (S19) respectively in randomly migrating RPE1 cells. The colocalization of PCDH7 with pMLC2 was pronounced in both leading **(Figure 1C, top)** and rear edges **(Figure 1C, bottom)**. We quantified the distribution of colocalization of PCDH7 and pMLC2 on the cell front and/or rear. Our results indicate that about 60 % of cells displayed only leading edge distribution of PCDH7 where it colocalized with p-MLC2. About 30% of cells displayed both rear and leading edge colocalization of PCDH7 and p-MLC2, whereas only rear edge presence of PCDH7 and p-MLC2 was very rare **(Supplementary Figure S1A)**.

### PCDH7 depletion affects cell migration and persistence

To investigate PCDH7’s role in cell migration, we used CRISPR/Cas9 method to knockout PCDH7 in RPE1 cells. According to the western-blotting analysis, PCDH7 expression is absent in PCDH7 knockout (KO) (**Figure 2A**). In addition to KO, add-back experiments were performed by the ectopic expression of PCDH7c to eliminate the possibility of off-target effects (Figure 2A). To analyze the effect of PCDH7 knockout on random migration, we performed single-cell tracking of RPE1 cells. Single-cell trajectories of KO cells were remarkably confined as compared to those of non-targeting (NT) control, whereas upon replenishment of PCDH7c (KO+PCDH7c), cells once again exhibited longer trajectories **(Figure 2B, Video 4)**. In addition, knockout cells displayed a loss of persistence phenotype (**Figure 2C** **and Video 4)**, which was recovered after the expression of PCDH7c in knockout cells (KO+PCDH7c) (**Figure 2C**). We also calculated angular displacement which is defined as the angular shift between two subsequent trajectories of a single cell. A higher proportion of small-angle changes (0^0^-36^0^) would indicate a stronger decisive behavior to migrate in the direction of initial migration and thus, a more persistent migration. Our analysis of angular displacements indicated a reduction in the proportion of the small-angle changes in KO cells compared to NT ones (**Figure 2D**). Upon recovery of PCDH7 expression (KO+PCDH7c), the proportion of small angle changes was increased (**Figure 2D**). We also used the shorter isoform, PCDH7b, for assessing the recovery upon knockout and observed a significant recovery in persistency (**Figure 2E**). Our data suggested that PCDH7 is required for persistent directional cell migration, and cells lose their directionality and thus, lose their persistent migration ability in PCDH7 depleted cells. Overall, PCDH7 KO cells displayed significantly reduced migration velocity than control cells. Ectopic expression of either isoform b or c recovered the velocity of migrating cells in knockout cells (**Figure 2F**).

**Figure 2.**
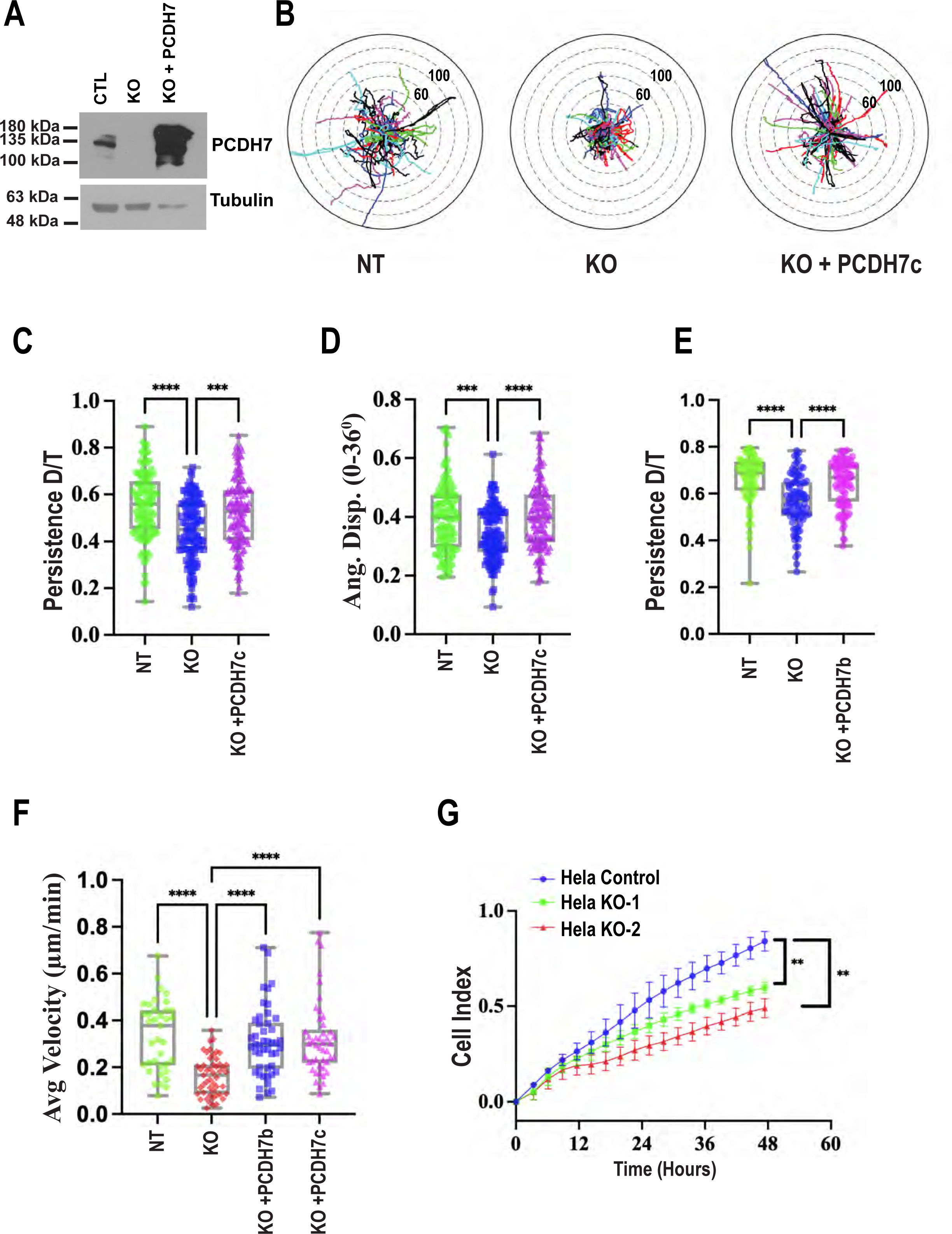
Perturbation of PCDH7 expression affects cell migration persistence and directionality. **(A)** Immunoblot analysis of PCDH7 using anti-PCDH7 antibody from CRISPR based PCDH7 knockout RPE1 cells and the recovered expression of PCDH7c in knockout cells (top). Tubulin is used as a loading control (bottom). **(B)** Representative trajectories from randomly migrating RPE1 cell populations of non-targeting (NT), knockout (KO), and add-back of PCDH7 (KO+PCDH7c). Concentric distance radial markers were placed at 20 µm apart. Measurement of migration parameters upon knockout (KO) and KO+PCDH7c for phenotype recovery: **(C)** Persistence as a ratio of direct distance and total distance (D/T), **(D)** measures of lower angle displacements (0-36°) n: NT=154, KO=174, KO+PCDH7c =195. **(E)** Persistence (D/T) for NT, KO, and KO+PCDH7b n: NT=85, KO=93, KO+P7B=96. **(F)** Comparison of average cell velocities in NT, KO, and recovery (KO+PCDH7b or KO+PCDH7c) cells. n: NT=37, KO= 49, KO+PCDH7b =46, KO+PCDH7c =46. One-way ANOVA was used for statistical analysis. **(G)** Real-time directed cell migration assay using xCELLigence RTCA for HeLaS3 cells. Blue circles indicate migration of HeLaS3 control cells whereas red squares and green triangles indicate migration behavior of two HeLaS3 knockout cells (HeLa KO-1 and HeLa KO-2, derived from two different sgRNAs) for PCDH7. Unpaired t-test was used for statistical analysis, *p<0.05, **p<0.01, ***p<0.001, ****p<0.0001.

To elaborate on the effects of PCDH7 in the context of cancer cells, we tested for migration defects in HeLaS3 cells. Two HeLa S3 PCDH7 knockout cell lines were generated using CRISPR/Cas9 system (Supplementary Figure S1B) and were used to compare their migration ability using xCELLigence Real-Time Cell Analysis for cell migration (Figure 2G). Essentially, this assay measures directional migration rather than random migration. Knockout cells displayed a significant reduction in cell migration compared to control cell lines. This result suggests that just like random migration, PCDH7 perturbation also affects directional migration.

### PCDH7 expression correlates with phosphorylation of MLC2

The role of PCDH7 in cell migration, and its colocalization with actomyosin, prompted us to explore its functional association with myosin II. To test whether PCDH7 plays a role in the activation of myosin II during migration, we biochemically tested pMLC2 levels (S19) in the non-targeting (NT) control and PCDH7 knockout (KO) HeLa S3 cells by Western Blotting. Strikingly, PCDH7 knockout HeLa S3 cells exhibited significantly reduced pMLC2 levels compared to control (**Figure 3A**). To examine the myosin activation spatially, we performed immunofluorescence for p-MLC2 levels in PCDH7 knockout RPE1 cells. We observed a dramatic reduction in pMLC2 levels at the leading and rear edges in PCDH7 knockout cells (**Figure 3B-D**). Upon expression of PCDH7 isoforms b or c in PCDH7 KO cells, p-MLC2 levels were significantly elevated in the leading and rear edges (**Figure 3B-D**). Our results suggest that PCDH7 expression levels alter MLC2 phosphorylation levels at the leading and rear edges of polarized cells.

**Figure 3.**
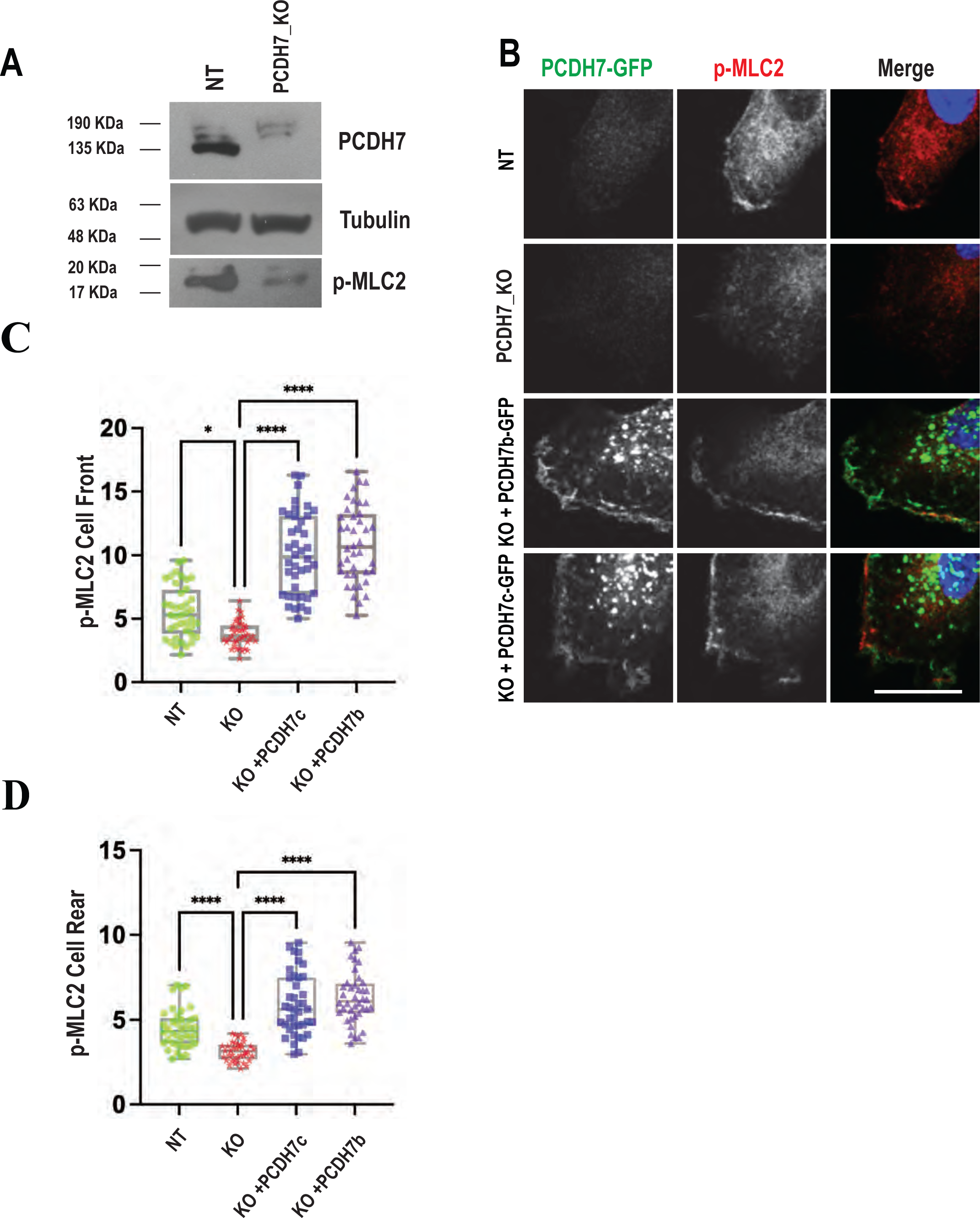
PCDH7 enhances myosin regulatory chain phosphorylation. **(A)** Immunoblotting analysis of PCDH7, pMLC2(S19) levels using anti-PCDH7 and anti-pMLC2(S19) antibodies in control (NT) and PCDH7-KO HeLaS3 cells. Tubulin is used as a loading control. **(B)** Representative images of PCDH7-GFP (α−GFP, green) and pMLC2(S19) (red) and DNA staining (DAPI, blue) in NT, KO RPE1 cells and recovery cells that express PCDH7-GFP isoforms (KO+PCDH7b, KO+PCDH7c). scale bars 20 µm. **(C)** Quantification of the pMLC2(S19) enrichment on cell front in NT (n = 40), KO (n = 40) and recovery (KO+PCDH7b (n = 40), KO+PCDH7c (n = 40)). **(D)** Quantification of the pMLC2(S19) enrichment on cell rear in NT (n = 40), KO (n = 40) and recovery (KO+PCDH7b (n = 40), KO+PCDH7c (n = 40)) One-way ANOVA was used for statistical analysis. *p<0.05, ****p<0.0001

### PCDH7 interacts with Myosin Phosphatase Target Subunit 1 (MYPT1)

Next, we focused on the molecular details through which PCDH7 expression affects MLC2 phosphorylation. Firstly, we performed pulldown of PCDH7c-GFP fusion protein followed by LC-MS/MS for identification of its interaction partners. Our results revealed components of actomyosin as the most significantly enriched proteins associated with PCDH7, including actin isoforms, subunits of myosin II isoforms **(Figure 4A, green box)**, ERM complex **(Figure 4A, blue box)** and various other actin cytoskeleton regulatory proteins. We observed clusters involved in the regulation of actin cytoskeleton, cadherin binding, and actomyosin regulation, among others **(Figure 4A, Supplementary Table S1)**.

**Figure 4.**
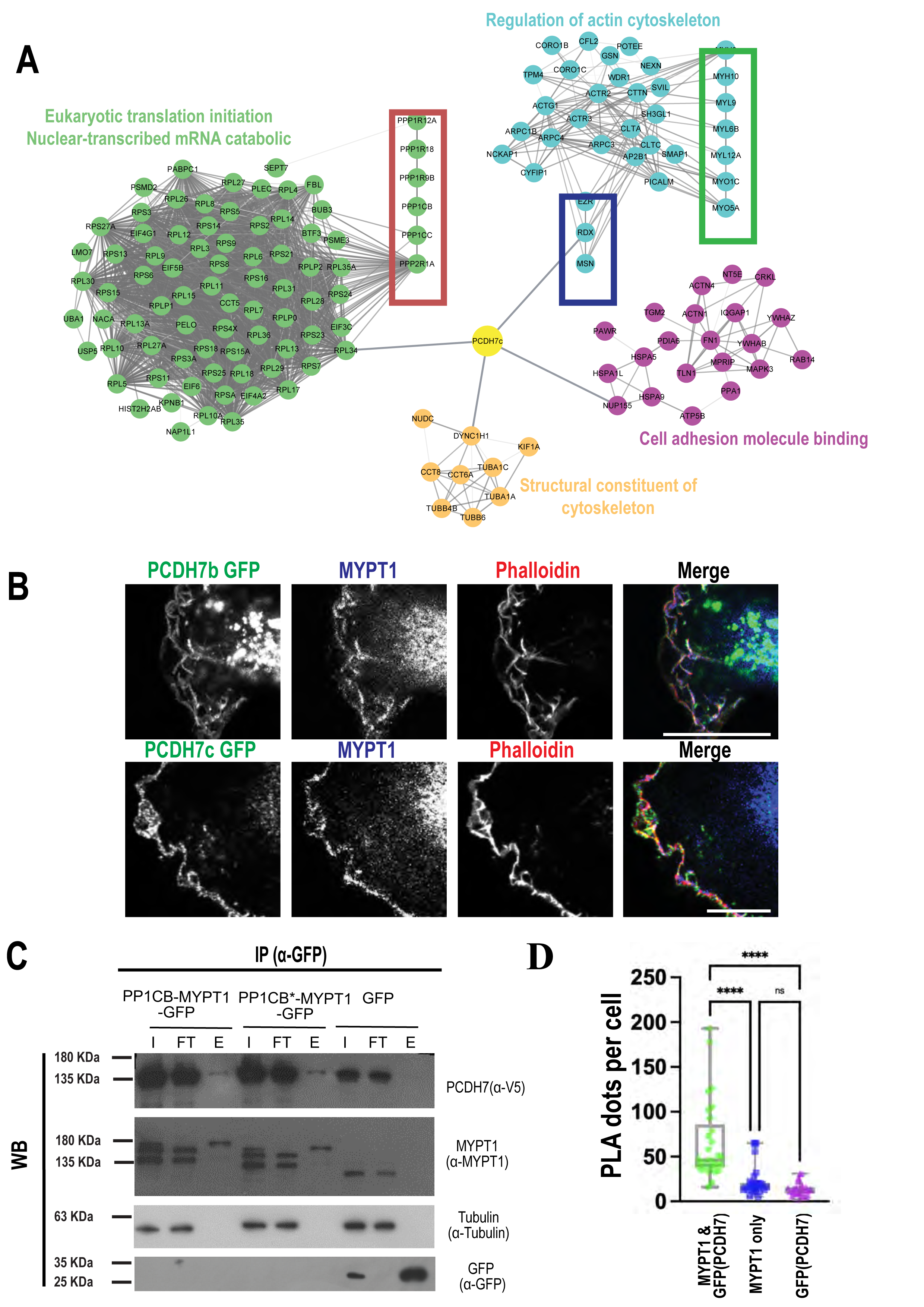
PCDH7 interacts with myosin phosphatase subunits. **(A)** Depiction of relevant clusters of PCDH7 isoform-c interactome determined by GFP mediated affinity pulldown followed by LC-MS/MS. Red inset depicts a set of phosphatase subunits and phosphatase regulatory proteins, green inset highlighting components of myosin complexes, and blue inset showing members of ERM proteins. **(B)** Representative images of RPE1 cells expressing PCDH7b- (top), PCDHc-GFP (bottom) stained with α-GFP (green), α-MYPT1 (red) and fluorescent phalloidin (blue). scale bars 20 µm **(C)** Immunoblotting analysis of GFP pulldown from GFP-PP1CB-MYPT1 fusion, GFP-PP1CB*-MYPT1 (D63A) mutant or only GFP (control) expressing Hela S3 cells with cotransfection of PCDH7c-V5 construct. Membranes were blotted using α -V5, α-MYPT1, α-GFP and α-tubulin antibodies. I (Input), FT(Flow Through), E (Elution) **(D)** Quantification of proximity ligation assay (PLA) showing fluorescence dots per cell for PCDH7-GFP (anti-GFP) and MYPT1 (anti-MYPT1) interaction (n=29) along with single primary antibody controls, MYPT1 (n=30) and GFP (n=30). One-way ANOVA was used for statistical analysis, ****p<0.0001.

Interestingly, we observed several component subunits of phosphatases such as MYPT1 (PPP1R12A), PP1cβ/δ (PPP1CB), PP1cγ (PPP1CC), and PPP2R1A, as interaction partners **(Figure 4A, red box)**. Both MYPT1 and PP1cβ are components of myosin phosphatase which regulates the phosphorylation of MLC2 (Kiss et al., 2019). The component PP1cβ is the catalytic subunit whereas MYPT1 is the myosin targeting subunit of myosin phosphatase holoenzyme (Shirazi et al., 1994). We performed immunofluorescence for MYPT1 and PCDH7 isoforms. Our results showed colocalization of both isoforms (PCDH7b and PCDH7c) with MYPT1 on the cell front (**Figure 4B**). To further analyze the interaction of PCDH7 and MYPT1, we performed cotransfection of PCDH7-V5 along with EGFP-PP1cβ-MYPT1, or EGFP-PP1cβ* (D63A)-MYPT1 (with catalytically inactive PP1cβ). Earlier work has shown a better substrate identification with mutant PP1 subunits (PP1cβ* (D63A)) wherein substrates are trapped in the complex during the PP1 catalytic activity (Wu et al., 2018). Our results showed the interaction of PCDH7c with both GFP-PP1cβ-MYPT1, and GFP-PP1cβ*(D63A)-MYPT1 (**Figure 4C**).

We also performed Proximity Ligation Assay (PLA) to further ascertain an interaction between PCDH7 and MYPT1. We used an anti-GFP antibody targeting PCDH7c-GFP and an anti-MYPT1 antibody for PLA. Our results indicated an interaction between PCDH7c and MYPT1 when anti-GFP and anti-MYPT1 are used together as compared to anti-GFP or anti-MYPT1 alone, further strengthening the evidence of interaction of PCDH7 and MYPT1 **(Supplementary Figure S2**, **Figure 4D**).

To test the interdependency between MYPT1 and PCDH7 enrichment, we performed immunofluorescence imaging of MYPT1 in PCDH7 knockout cells. We did not observe a significant change in MYPT1 levels at the leading edges upon PCDH7 KO, however ectopic expression of PCDH7c or PCDH7b in KO cells significantly recruited more MYPT1 to the leading edges **(Supplementary Figure S3A,B)**. Taken together, our results showed that PCDH7 interacts with myosin phosphatase subunit MYPT1 and that the PCDH7 expression correlates with MYPT1 enrichment at the leading edge.

### PCDH7 facilitates Ezrin Activation at the Leading Edges

All the components of the ERM complex were identified as the interactor of PCDH7 in our mass spectrometry analysis (**Figure 4A**), which prompted us to investigate the relation between PCDH7 and the ERM (ezrin/radixin/moesin) proteins. The activation of the Ezrin protein is triggered by the phosphorylation of its carboxyl-terminal (T567), which stimulates its F-actin binding (Bosk et al., 2011). The active form of Ezrin is crucial for the establishment of leading edge and cell motility (Prag et al., 2007). Given the fact that Myosin Phosphatase regulates ERM proteins by dephosphorylating them (Fukata et al., 1998; Kiss et al., 2019), we tested whether PCDH7 affects ezrin activation in RPE cells. We compared phospho-ERM levels at the leading and rear edges in control, PCDH7 KO and recovery cells (PCDH7 KO+PCDH7c and PCDH7 KO+PCDH7b) by performing immunofluorescent analysis **(Figure S4A)**. In PCDH7 knockout, p-ERM levels are reduced at the leading and rear edges and this reduction gets replenished upon PCDH7b or PCDH7c expression **(Supplementary Figure S4B, C)**.

### PCDH7 Expression Correlates with Inhibitory MYPT1 Phosphorylation and Chemical Inhibition of Phosphatase Activity Recovers PCDH7 Knockout Phenotypes

Our results suggest that the absence of PCDH7 affects the p-MLC2 and p-ERM levels at the front and rear edges of migratory cells. Given the interaction between PCDH7 and MYPT1, we hypothesized that PCDH7 might have an inhibitory effect on MYPT1. To test the inhibitory effect of PCDH7 expression on MYPT1, we monitored phosphorylation of an inhibitory site on MYPT1 (Thr696) upon ectopic expression of PCDH7. Strikingly, the levels of phospho-MYPT1(Thr696) significantly increased by the PCDH7 expression **(Figure 5A, B)**.

**Figure 5.**
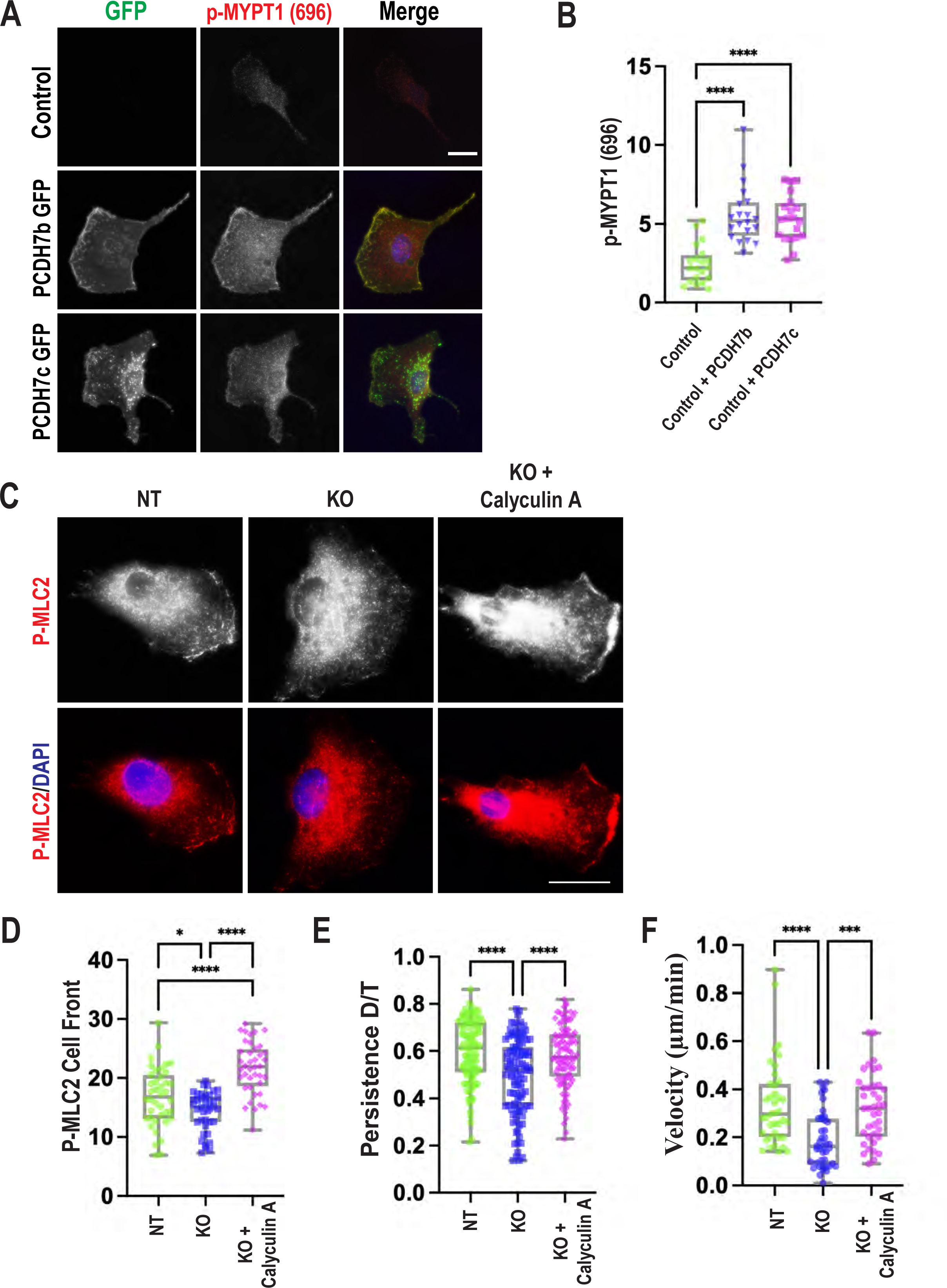
Ectopic expression of PCDH7 increases inhibitory phosphorylation of MYPT1 (Thr696) whereas chemical inhibition of phosphatase activity recovers reduced pMLC2 levels and persistence of migration in KO cells. **(A)**. Representative images of WT, PCDH7b GFP and PCDH7c GFP expressing RPE1 cells stained with α-GFP and α-pMYPT1(696) and Hoechst. scale bar 20 μm. **(B)** Quantification of p-MYPT1 on the cell front in control (N=20), upon ectopic expression of PCDH7 isoforms (PCDH7b GFP (N=20), PCDH7c GFP (N=20)). **(C)** Immunofluorescence images showing NT, KO, and Calyculin A (15 pM) treated KO (KO + Calyculin A) RPE1 cells stained with α-pMLC2 (S19) and DAPI. Scale bar 20 µm **(D)** Quantitative comparison of pMLC2(S19) levels on cell front in NT (n=40), KO (n=40), and KO + Calyculin A (n=40). **(E)** Comparison of persistence of migration in NT (n=102), KO (n=105), and KO+ Calyculin A (n=108) (Video 5). **(F)** Comparison of velocities of NT, KO, and KO+calyculin A, NT (n=39), KO (n=42), and KO+ Calyculin A (n=39). One-way ANOVA was used for statistical analysis, p value: *p<0.05, **p<0.01, ***p<0.001, ****p<0.0001.

To test the phosphatase activity-dependent role of PCDH7, we employed calyculin-A, a potent chemical inhibitor of protein phosphatase activity which is more selective against PP1 than PP2 (Kita et al., 2002). We hypothesized that the enhanced activity of MYPT1 at the cell front can be controlled by limited inhibition of phosphatase activity in the PCDH7 knockout cells which would recover the phenotypes of reduced p-MLC2 levels and cell motility. Although the IC50 of calyculin-A for PP1 lies in the nanomolar range (Leung et al., 2002; Tanaka et al., 2007), RPE1 cells exhibited rounding behavior and subsequent detachment of the cells in nanomolar concentrations (data not shown). Instead, we used the calyculin-A at a picomolar concentration where cell morphologies and migration behavior were not hampered. Indeed, we observed that at 15 pM concentration, myosin phosphorylation levels at the leading edges were significantly enhanced compared to the knockout **(Figure 5C, D)**. We also checked calyculin-A treatment dependent recovery of cell migration defects (both persistency and velocity phenotypes) in knockout cells. In 15 pM calyculin-A treated RPE1 cells, both persistence and velocity of PCDH7 KO cells are significantly recovered **(Video 5 and** **Figure 5E-F**). Thus, the chemical inhibition of phosphatase activity aids in the recovery of p-Myosin levels and migration persistency which further strengthens the link between PCDH7/Myosin-Phosphatase/Myosin axis and cell motility.

### *In vivo* radial neuronal migration is enhanced by PCDH7 overexpression

Protocadherins are highly expressed in neurons and an increasing body of evidences show crucial roles of protocadherins in the brain (Hayashi and Takeichi, 2015). PCDH7 was previously named BH-PCDH (brain and heart protocadherin) owing to its high expression in these tissues (Yoshida et al., 1999). After observing the role of PCDH7 in cultured cells, we aimed to uncover the role of PCDH7 in cell migration *in vivo*. To this end, we studied the effect of PCDH7 expression during neuronal migration in the cerebral cortex. We used *in utero* electroporation and were able to overexpress PCDH7b in ventricular neuronal progenitor cells of E14.5 (**Figure 6A**). We also expressed GFP marker (pCAGGS_IRES_GFP) to observe the migration of the transfected neurons at 3 days post electroporation (E17.5). In the control (EGFPN1) samples, we observed expected migration of neurons, 40% of them were able to reach the cortical plate (CP), 50% of transfected cells migrated along the subventricular zone-intermediate zone (SVZ-IZ) and 10% of them were located to the ventricular zone (VZ) (**Figure 6B**). Upon overexpression of PCDH7b in the mouse embryonic cortex, we observed a significant increase in the proportion of electroporated cells that migrated to the CP. Cells that expressed PCDH7b had an increased rate of migration, hence PCDH7 GFP-positive neurons exited the VZ and migrated toward the CP **(Figure 6A, lower panel and** **Figure 6B**). This result indicates that overexpression of PCDH7 in the subventricular neuronal migratory population enhances the migratory behavior of newly born neurons.

**Figure 6:**
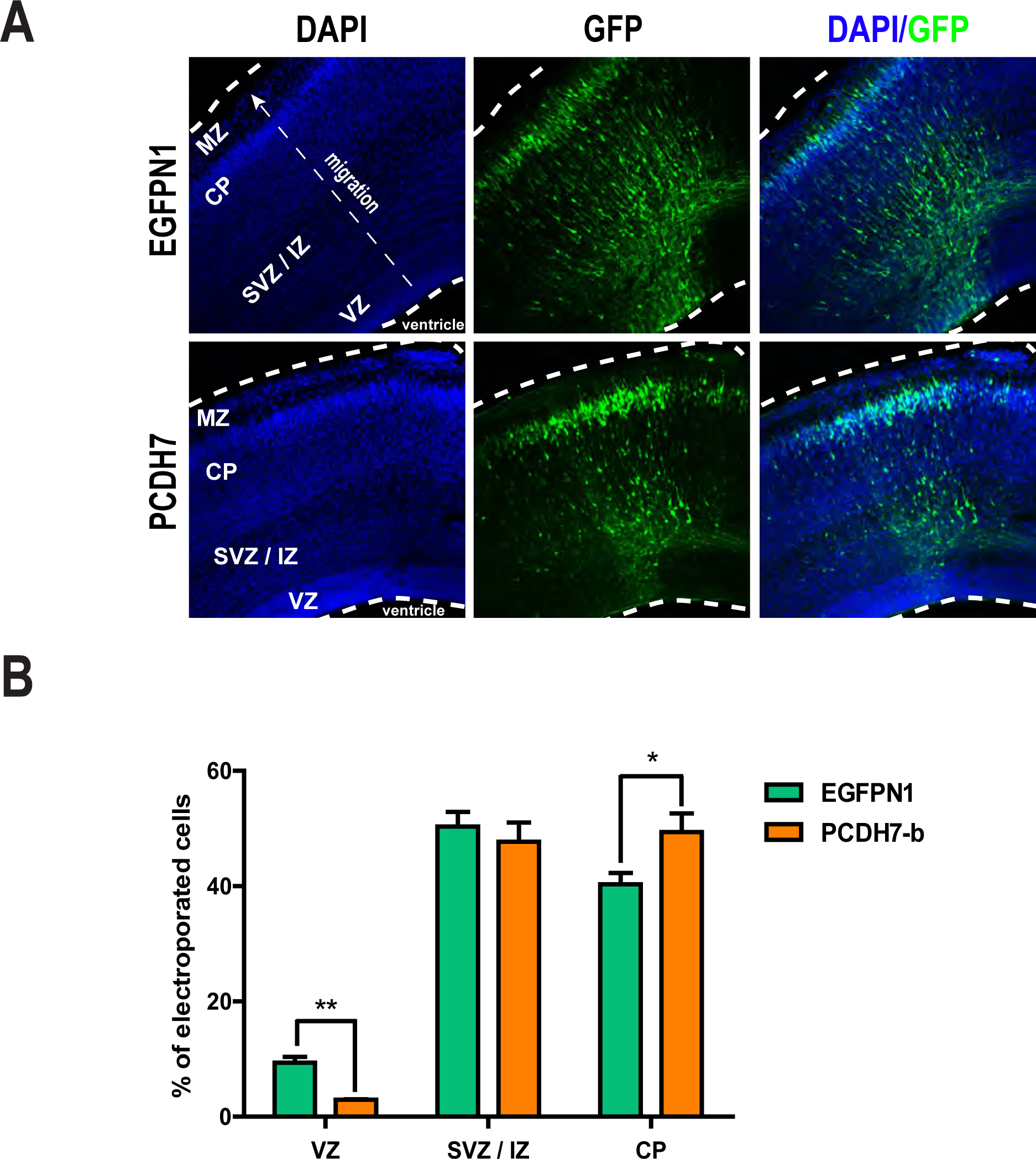
PCDH7 overexpression enhances radial neuronal migration in mouse developing cerebral cortex. **(A)** Immunofluorescence images of E17.5 coronal cryosections *in utero* electroporated with EGFPN1 (control) or PCDH7b-EGFPN1 and pCAGGS_IRES_GFP at E14.5. GFP immunostaining was performed to visualize migration pattern of transfected cells and DAPI was used to stain nucleus. VZ, ventricular zone; SVZ-IZ, subventricular zone-intermediate zone; CP, cortical plate; MZ, marginal zone. Scale bar, 100 μm**. (B)** Quantification of *in utero* electroporation experiment of PCDH7b. Cortical sections were divided into zones and normalized percentages of GFP-positive neurons counted in each zone are plotted as bar graphs. (For PCDH7b; a total of n=3373 for EGFPN1, n=3316 from 2 biological replicates were included. Lines represent S.E.M. Unpaired two-tailed *t* test, p value: *p<0.05, **p<0.01.

## Discussion

Despite its importance, how the cortex is organized and communicates with the plasma membrane during cell migration is not fully understood. Crosslinked actin network, myosin motors and the associated proteins make up the cortex which lies right under the plasma membrane. The plasticity of the cortex allows the reversible association of the cortex with the plasma membrane and helps in the mechanical transduction of motile cells (Salbreux et al., 2012; Svitkina, 2018). This is indispensable for every step of cell migration: breaking of cell symmetry, protrusion of the leading edge, adhesion to the substrate, contraction and detachment of the rear edge (Petrie et al., 2009). High turn-over of proteins and dynamic interactions in the cortex are thought to contribute to the cortical plasticity; however, its molecular details remain to be determined.

In the present work, we showed that a member of the cadherin superfamily, namely protocadherin-7 (PCDH7), displays polarized behavior in a migrating cell with accumulation at the cell leading and rear edges. In a directed migration, cell persistence is defined as the ability of straight-line motion which is correlated with the migration speed through maintenance of cell polarity by actin flow (Maiuri et al., 2015). We showed that the perturbation of PCDH7 through CRISPR/Cas9 based knockout reduces the persistence and the velocity of migrating cells. We further investigated that PCDH7’s perturbation reduces phosphorylation of MLC2 and its overexpression enhances phosphorylation of MLC2 at the leading and rear edges. Myosin motor activity is a highly regulated process that is reversibly controlled by phosphorylation/dephosphorylation of the MLC2 through activities of Rho-Kinase, Myosin Light Chain Kinase and Myosin Phosphatase respectively (Leung et al., 2002; Somlyo and Somlyo, 1994). We suggest that PCDH7 involves in the regulation of activity of myosin at the leading and rear edge to cell polarity and enhances cell persistence. In line with this conclusion, our previous studies demonstrated that PCDH7 functions in building up rounding pressure, which is another myosin II dependent function (Ozlu et al., 2015). In addition, PCDH7 inhibits homotypic cell-in-cell structure by increasing myosin phosphorylation (Wang et al., 2020).

How does PCDH7 regulate myosin activity during cell migration? We showed that PCDH7 forms a complex with MYPT1, and PP1cβ which are subunits of myosin phosphatase. MYPT1 is responsible for directing the complex to myosin (MLC2), and PP1cβ is the catalytic subunit that carries out myosin dephosphorylation (Kiss et al., 2019). We confirmed the interaction between PCDH7 and MYPT1 through co-immunoprecipitation, immunofluorescence and Proximity Ligation Assay. We also showed that MYPT1 enrichment on the cell front and the cell rear have a direct correlation with PCDH7 expression; its enrichment is reduced upon knockout and is enhanced upon overexpression. Importantly, the inhibitory proteo-form of MYT1 also accumulated at front edges upon PCDH7 ectopic expression. PP1α, which is an isoform of PP1cβ, has previously been shown to interact with PCDH7 through the canonical PP1α interaction motif (RRVTF) in its cytoplasmic domain (CM3) (Wang et al., 2020; Yoshida et al., 1999). The myosin phosphatase targeting subunit, MYPT1 directly binds to PP1cβ by its RVxF motif (Kiss et al., 2019). Interestingly, in addition to the longer isoform (PCDH7c), the short version PCDH7b, which does not harbor (RRVTF) motif, also colocalized with myosin phosphatase and rescued the migration phenotype. Therefore, we propose that PCDH7 and myosin phosphatase interaction is mainly through their direct interactions with MYPT1 subunit in addition to the PP1cβ-interaction.

In addition to myosin, our study suggested that another essential component of cell migration, the ERM complex (Yu et al., 2004) is also regulated by PCDH7. Like MLC2, ERM is also a substrate of Myosin Phosphatase (Kiss et al., 2019). Our analysis demonstrated that PCDH7 interacts with ERM and facilitates the recruitment of phosphorylated (activated) Ezrin to the leading edges. The expression of PCDH7 correlated with the enhanced phosphorylation level of ERM at the migratory cortex. We propose that MYPT1-PCDH7 interaction contributes to the localized activation of ERM at the migratory cortex. Thus, through the PCDH7-MYPT1-ERM axis, PCDH7 may act as a component of the connecting link between the plasma membrane and the actin cytoskeleton which contributes to the establishment of the polarized cortex during migration. We observed that PCDH7 overexpression led to an increase in phosphorylation dependent inhibition of MYPT1. In addition, chemical inhibition of phosphatase activity using Calyculin A increased MLC2 phosphorylation level and rescued PCDH7 migration phenotype. It has been previously shown that PCDH7 suppresses PP2A activity by mediating inhibitory interaction with SET, an inhibitor of PP2A (Zhou et al., 2017). Similarly, PCDH7 may not directly inhibit myosin phosphatase activity but rather suppresses its activity by promoting inhibitory interactions. We also cannot exclude the possibility that PCDH7 suppresses myosin phosphatase activity by influencing Rho-ROCK modulators. A recent study reported that PCDH7 involves in the activation of Rho signaling during osteoclast differentiation (Kim et al., 2021). Further study is necessary to elucidate the molecular details of PCDH7 dependent regulation of myosin phosphatase activity.

Recent work showed that PCDH7 determines the morphology of the dendritic spine (Wang et al., 2020). It is plausible that PCDH7 shape the dendritic morphologies or that its orthologues function in retinal axon growth and guidance (Heggem and Bradley, 2003; Leung et al., 2015; Leung et al., 2013; Piper et al., 2008) via a similar mechanism, namely by regulating phosphorylation dynamics of MLC2 (Hodges et al., 2011; Rex et al., 2010; Zhang et al., 2005) and ERM. The roles of cadherins in collective cell migration have been increasingly investigated (Theveneau and Mayor, 2012). OL-pc/PCDH10, a member of the δ-2 protocadherin group has been shown to mediate cell migration when cells are in contact with other cells but not singly (Nakao et al., 2008). Like other members of the cadherin family, PCDH7 is also enriched at cell-cell junctions. As Myosin IIA and Myosin IIB isoforms are involved in adherent junctions integrity and biogenesis (Heuze et al., 2019), PCDH7’s regulation of myosin activity at cell-cell junctions might contribute to the collective cell migration. More work is required to decipher the role of PCDH7 in collective cell migration. Yet, our study supports the emerging role of protocadherins in migration dependent physiological and pathological processes.

To conclude, we have shown that PCDH7 is an important regulator of directed migration where it displays a polarized localization at the leading and rear edge and spatially regulates myosin and ERM activity. We propose that PCDH7 imparts this regulation through its interaction with myosin phosphatase at the cell cortex. Here we propose a new component of cell migration that plays an important role in the plasticity of the cell cortex. Further investigation of the molecular mechanisms behind the interaction of PCDH7, myosin phosphatase and myosin will provide important clues about the polarization and the plasticity of the cell migration machinery. It will also shed new light on the signaling pathways behind metastasis.

## Materials and Methods

### Plasmids and Lentiviral Transduction

Following vectors are used for cloning b and c isoforms of PCDH7: eGFP-N1, pLenti CMV GFP Puro (addgene#17448), PLEX_307 (Addgene#41392). eGFP-PP1CB-MYPT1 and eGFP-PP1CB(D63A)-MYPT1 were provided by Mathieu Bollen from KU, Leuven and generated as previously described (Wu et al., 2018). Lentiviral constructs expressing Lifeact-RFP, Ezrin-RFP, and Cortactin-RFP were provided by Michael Way from Francis Crick Institute, UK. To produce lentiviruses, HEK293T cells are transfected with the following vectors psPAx2 (Addgene #12260) and pMD2.G(Addgene # 12259) using Lipofectamine 2000. The supernatants were collected at 48-72 hours. RPE1 cells were infected with Lentiviral packaged particles by supplementing 8 μg/ml protamine sulfate (P4505; Sigma-Aldrich), the virus infected cells were selected by overnight puromycin (7 μg/ml) treatment.

### CRISPR/Cas9 Mediated Knockouts of PCDH7

Guide RNA sequences for PCDH7 were selected using mit.crispr.edusoftware. For generating HelaS3 knockout cell lines, following sequences 5’-CGACGTCCGCATCGGCAACG-3’, and 5’-CATCGTGACCGGATCGGGTG-3’ were used separately as guide RNAs. For generating RPE1 knockout cell lines, a targeting guide with sequence 5’-TTGCCGATGCGGACGTCGGC – 3’ was used. For non-targeting control, the following guide sequence 5’-ACGGAGGCTAAGCGTCGCAA - 3’ was used. The guides were cloned into pLenti-CRISPRv2 using BsmBI restriction digestion of the backbone using recommended protocols from Zhang lab (Sanjana et al., 2014). HeLa S3 and RPE1 cells were transiently transfected with plentiCRISPRv2 plasmid containing PCDH7 guides and non-targeting control. Monoclonal cells were prepared with dilution and selected for the absence of PCDH7 expression using immunoblotting.

### Live Imaging and Single-cell tracking Analysis

To characterize the migration phenotype by using single-cell tracking method, RPE1 CRISPR knockout and control cells expressing GFP fluorescent protein were plated in a diluted manner, after 12 hours of incubation in DMEM/F12 supplemented with 10% FBS and 1% Penicillin/Streptomycin. To image migrating cells, they were placed at humidified chamber at 37 °C and 5% CO_2_ during the data acquisition. OlympusPro IX71 equipped with Andor iXon EMCCD camera and Leica DMi8 live imaging microscopes with 10x air objective lens were used for time-lapse imaging of cells. All videos were recorded for several hours with a frame rate of 10 minutes. A custom single-cell tracking and motility characterization code written in MATLAB (R2020b, The MathWorks) were partially adapted from previous studies (Conkar et al., 2019; Seker et al., 2019). Briefly, tracking analysis composed of four main sections includes video loading, image smoothing, segmentation and cell linkage steps respectively. First, low-pass filter was used to remove pixel noise from background subtracted raw videos. 2D Gaussian filter with an average radius of 2-5 pixel was set to smooth images. After computing the intensity distribution, threshold filter was applied to separate cells. To group pixels belonging to each candidate cell body, an array of 8-neighbourhood for each pixel was used to compute the connectivity. The centroid position of labelled cells was identified by using local maximum detection. Finally, a list of coordinates from each frame was analyzed by tracking algorithm to find the linked cells in consecutive frames. Nearest neighbor method was applied to determine most plausible matches of cells. An average distance of 40 pixels was set as a cutoff value of displacement. After computing Euclidian distance between frames, trajectories were identified by minimizing the distance function of matched cells. The positions of linked cells were finally saved to compute motility dynamics. Supplementary Figure S5 represents an example of cell tracking from a live cell imaging. The persistence defined as the ratio of direct (D) to total (T) distance was used as an indicator of directed motion. Cells moving linearly have a value approaches to 1 while the persistence ratio decreases as they tend to move randomly. An average of angular displacement (θ) indicating the rotational movement of cells was computed by using the following formula,

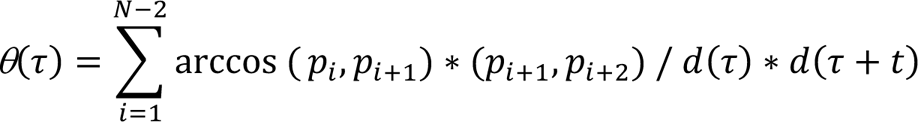

where θ(*τ*) denotes the degree between three consecutive frames. *P*_i_ and N represent X-Y position vector and the total number of video frames respectively. d(τ) *and d*(τ + *t*) are the mean squared distance between two frames. *t* denotes the time resolution. To compute the average velocity, total distance over time was calculated by using the cell coordinates that were computed at the tracking section.

### xCELLigence Real Time Cell Migration experiment

Real time cell migration was performed as recommended using xCELLigence RTCA-DP (ACEA Biosciences Inc., CA, USA). Briefly, 10% FBS containing DMEM was used at the bottom and 1% FBS DMEM was inserted at the top to create downward chemoattractant gradient. 30,000 cells per cell line were plated on the top chamber in 100 µl of medium and let set for 1 hour for the cells to attach to the surface of the top plate. 16 well cell invasion and migration (CIM) plates were used for this experiment. RTCA migration study was performed for 48 hours.

### GFP Pull-down for Protein Interaction Analysis

The cells were harvested in HBSS solution (Diagnovum D408) and lysed in a buffer [pH 7.5 10mM Tris, 150 mM NaCl, 0.5% NP40, 1.26 mM CaCl2, 0.5 mM MgCl2.6H2O, 0.4 mM MgSO4.7H2O] supplemented with protease inhibitor (complete^™^, Mini, EDTA-free Protease Inhibitor Cocktail REF 18836170001) and phosSTOP phosphatase inhibitor cocktail (Roche 4906845001)] at 4^0^C for 30 minutes before clearing lysate by centrifugation at 4^0^C. GFP nanotrap agarose beads (Chromotek gta) were used for pulldown of GFP and GFP fusion proteins as manufacturer’s instructions. Samples from input, flow through and elution were loaded to the SDS-PAGE.

### Protein Identification by Mass Spectrometry

On-bead tryptic digestion was performed. Briefly, the protein bound agarose beads were washed with 50mM Ammonium bicarbonate followed by reduction with 100mM DTT and alkylation with 100 mM iodoacetamide. MS Grade Trypsin (Pierce) is used for the digestion followed by desalting using C8 Stagetips. The resulting peptides were analyzed in LC-MS/MS (Q Exactive, Thermo Fisher Scientific). Proteome Discoverer (1.4) is used for database search. Spectral counts of identified proteins were used to calculate the Bayesian false discovery rate (BFDR) and the probability of possible protein-protein interactions by SAINTexpress v.3.6.3 with -L option. Proteins were filtered out according to the SAINTscore (>0.5) and BFDR value (<0.05). Significant protein hits with 1 SAINTscore were loaded into the STRING database v11.0 (Szklarczyk et al., 2019) by Cytoscape StringApp (Doncheva et al., 2019) with 0.7 confidence. MCL clustering (inflation value: 2.5) of the network was performed by Cytoscape (version 3.7.2) and its plugin clustermaker (Cline et al., 2007) GO and KEGG enrichment analysis of the network was performed via g:Profiler (Raudvere et al., 2019).

### Proximity Ligation Assay

Duolink Proximity Ligation Assay Kit (92101, Sigma-Aldrich) was used according to the manufacturer’s instructions. RPE1 cells were plated on crossbow shaped micropattern coverslips following the manufacturer’s instructions (10-950-10-18, CYTOO). Two control samples with only primary antibodies (anti-MYPT1, sc-514261, Santa Cruz; and anti-GFP, 2555S, Cell Signaling Technology), and one target sample with both primary antibodies were used. Cells were imaged in the Texas Red channel for quantification of PLA, DAPI for nucleus and FITC for PCDH7 expression. Images were processed in ImageJ for finding maxima to highlight the particles and for calculating the number of PLA dots. Datasets were plotted in PRISM GraphPad and statistical analysis was performed using one-way ANOVA.

### Immunofluorescence and Quantification

Cells were fixed with 3% paraformaldehyde at 37°C for 8 minutes, followed by 0.1% Triton-X 100 in PBS permeabilization .blocking was done in 2 % BSA 0.02% NaN3 in 1x PBS for 1 hour. Following antibodies are used in immunofluorescence: anti-GFP goat (gift from Hyman lab, 1:10,000), anti-GFP mouse (11814460001, Roche 1:1000), anti-MYPT1 mouse (sc-514261, Santa Cruz 1:200), anti-pERM rabbit (Cell Signaling Technology 3726s, 1:500), phalloidin iFlour555 (ab176756, Abcam), and phalloidin Alexa flour 647 (A22287, Thermo Fisher Scientific) and phospho-myosin light chain 2, S19 mouse (3675, Cell Signaling Technology, 1:500), phospho-MYPT1(Thr696)(PA5-38297, Sigma 1:100). DAPI or Hoechst were used for nuclear labeling. Mowiol was used for the mounting of coverslips. Imaging was performed by 63× Plan Apo 1.4 NA oil-immersion objective of Leica DMi8 wide-field microscope. Z stacks were acquired in 3µm and the maximum projection was performed before quantification. ImageJ software was used for quantification. Images were first converted to 8-bit version and then a region of interest was determined as a 64 pixel^2^ square. After five different areas were measured in the cell cortex for cell front and cell rear edges, their averages were calculated. Five different extracellular regions were measured as background signal and subtracted from cell front and cell rear signals.

### Immunoblotting

Anti-PCDH7 mouse (139274, Abcam) 1:200, anti-Tubulin (3873S, Cell Signaling Technology) 1:2000, anti-MYPT1 mouse (sc-514261, Santa Cruz) 1:500, anti-V5 mouse (Invitrogen R96025) 1:2500, anti-GFP antibodies (Cell Signaling Technology 2555S, and Santa Cruz Sc-8334) 1:1000, and anti-phospho-myosin light chain 2, S19 mouse (3675, Cell Signaling Technology) 1:500 were used. HRP labelled secondary antibodies at 1:2000 dilution were used, anti-mouse (7076 Cell Signaling Technology) and anti-rabbit (7074 Cell signaling technology).

### *In-utero* electroporation

*In utero* electroporation experiment was performed as previously described (Guzelsoy et al., 2019; Saito, 2006). *In utero* electroporation protocol was carried out using E14.5 pregnant CD1 mice. Briefly, pregnant mice were anesthetized with isoflurane and laparotomy was performed to expose the uterine horns. 500 ng of control (EGFPN1) or PCDH7b-EGFPN1 plasmids was mixed with 500 ng pCAGGS_IRES_GFP marker to observe the transfected cells and 0.1% Fast Green to monitor the injection (QIAGEN Endofree Plasmid Maxiprep kit were used to prepare all plasmids). Plasmid mixes were injected into lateral ventricle visible through the translucent uterine wall with pulled glass capillaries. After injection, plasmid DNA mix were transfected into the neuronal progenitor cells located in the VZ by electroporation with the settings 0.5 pulses, 30 V, 400 ms intervals using an electroporater (BTX Harvard Apparatus, ECM830) and 5 mm tweezertrodes (BTX Harvard Apparatus). Embryos were placed back into the body cavity and allowed to grow for three days. At E17.5, electroporated embryonic brains were collected and fixed with 4% paraformaldehyde overnight at 4°C. 50 µm coronal sections of the embryonic brains were prepared using cryostat (CM1950, Leica Biosystems). Sections were stained with anti-GFP (Aves Labs, Cat. #GFP-1020, 1:1000) antibody to look for the pattern of the migration. Nikon Eclipse 90i confocal microscope was used to capture optical z-series through 50 µm images and NIS-Elements AR software was used to generate maximum intensity projections. 10 bins grid was placed onto the images using Adobe Photoshop CC (Adobe Systems) and number of GFP-positive cells were counted for each bin with ImageJ (NIH) cell counter option (85). Data were transferred to Excel (Microsoft Office) for analysis. Bins 1-5 indicate CP, bins 6-9 indicate SVZ-IZ, and bin 10 indicates VZ. Graphs were plotted using GraphPad Prism 5 software. *In utero* electroporation protocol was performed in accordance with the guidelines for the care and use of laboratory animals of Koç University and the Ministries of Food, Agriculture and Livestock, Forestry and Water Management of Turkey. CD1 mice were accommodated, sustained, and bred at the Koç University Animal Facility. Ethics approval was received from Koç University Institutional Animal Care and Use Committee.

### Statistical Analysis

For the analysis of single cell tracking experiment, proximity ligation assay, and immunofluorescence quantification one-way ANOVA was used. For RTCA cell migration and *in-utero* electroporation, unpaired t-test was used. p-value -0.05 was used as cut-off for significance. Plots were generated using Graphpad PRISM software.

## Acknowledgement

We thank Koc University Proteomics Facility (KUPAM) for proteomics services and support. We thank imaging facility at Koc University (CMIC) for facilitating imaging experiments. We thank Dr. Mathieu Bollen from KU Leuven for supplying MYPT1 and PP1CB fusion vectors. We thank Dr. Michael Way from the Francis Crick Institute for providing lifeact-RFP, and Ezrin-RFP constructs. We thank Aydanur Senturk for her inputs and critical reading of the manuscript. This work is funded by TUBITAK 1001, 116Z305 to NO. H.B. was supported by ITU-BAP grant 2020-42579.

## Author Contribution

NO and MHQ conceived the project and designed the experiments. MHQ and MTC performed the experiments. EB assisted in the experiments. Analysis of single cell tracking for migration phenotype was performed by HB. Proteomics data analysis was performed by ABK. RTCA experiment was performed by ZCUK, in utero experiment was performed by CA. MHQ and NO prepared the manuscript.

## FIGURE LEGENDS

**Figure S1. Characterization of PCDH7 localization and depletion (A)** Distribution of PCDH7c and p-MLC2 colocalization on cells front and/or rear edges, N=29 (Cell number is normalized to 1). **(B)** Western blotting analysis of PCDH7 knockout in HeLaS3 cells using α-PCDH7 and α-tubulin antibodies. Two different guide RNAs were used for the generation of two different clones. PCDH7 depletion is confirmed and tubulin is used as loading control.

**Figure S2. PCDH7 and MYPT1 interaction by proximity ligation assay** Representative images from proximity ligation assay depicting fluorescent signal from interaction of MYPT1 and PCDH7-GFP (Red dots) using anti-GFP and MYPT1 antibodies (MYPT1+GFP, upper panel) as well as the control groups where only anti-MYPT1 (middle) or only anti-GFP (lower panels) antibodies are used. Scale bar 20 µm.

**Figure S3. PCDH7 expression correlates with the enrichment of the MYPT1 component of myosin phosphatase. (A)** Representative images of non-targeting (NT), knockout (KO), and KO+PCDH7c and KO+PCDH7b expressing RPE1 cells stained against anti-GFP and anti-MYPT1 antibodies. Scale bar 20 µm. **(B)** Quantification of MYPT1 enrichment on the cell front in NT (n=40), KO (n=40), KO+PCDH7c (n=40) and KO+PCDH7b (n=40) cells. One-way ANOVA was used for statistical analysis,**p <0.01, ***p<0.001, ****p<0.0001.

**Figure S4. PCDH7 expression correlates with ERM phosphorylation at the cell front and cell rear**. **(A)** Representative immunofluorescence images of NT, KO and KO+PCDH7c and KO+PCDH7b RPE1 cells stained against anti-GFP, anti-pERM and labeled phalloidin to visualize PCDH7-GFP (Green), p-ERM (Red), actin (Magenta). Scale bars 20 µm. (**B)** Quantification of p-ERM enrichment on the cell front in NT (n=40), KO (n=40), and in recovery KO+PCDH7c (n=40) and KO+PCDH7b (n=40). (**C)** Quantification of p-ERM enrichment on the cell rear in NT (n=40), KO (n=40), KO+PCDH7c (n=40) and KO+PCDH7b (n=40). One-way ANOVA was used for statistical analysis, **p <0.01, ****p<0.0001.

**Figure S5:** Trajectory analysis of cells to determine cell connectivity in consecutive frames. X-Y coordinates from tracking section was used to compute cell persistence and angular displacement. Raw videos were loaded for segmentation of image frames. **(A)** representative sequence of images from a video. **(B)** shows cell regions (color-coded) and centroids (white dots) determined after computing pixel connectivity. **(C)** Right panel shows cell trajectories (magenta lines) that were overlaid on raw images. To determine linked cells, nearest neighbor method was applied by minimizing mean square distance between cells. The positions of tracked cells were used to determine the cell dynamics. (Scale bar, 40 µm).

**Video 1**: Confocal scanning imaging of RPE1 cell expressing PCDH7c-GFP and lifeact-RFP. Individual frames were collected 10 minutes apart.

**Video 2**: Confocal scanning imaging of RPE1 cell expressing PCDH7c-GFP and lifeact-RFP. Individual frames were collected 10 minutes apart.

**Video 3**: Confocal scanning imaging of RPE1 cell expressing PCDH7b-GFP and lifeact-RFP. Individual frames were collected ∼2 seconds apart.

**Video 4**: Single cell tracking of randomly migrating NT control, PCDH7 KO, and KO+PCDH7c recovered RPE1 cells.

**Video 5**: Single cell tracking of NT (control), PCDH7 KO and Calyculin treated PCDH7 KO (+Calyculin) RPE1 cells.

**Supplementary Table 1.** Proteins identified as interactors upon PCDH7c-GFP pulldown followed by LC-MS/MS.

