## Supplementary Figures for "PCDH7 Promotes Cell Migration by Regulating Myosin Activity"

Qureshi et. al., Figure S1

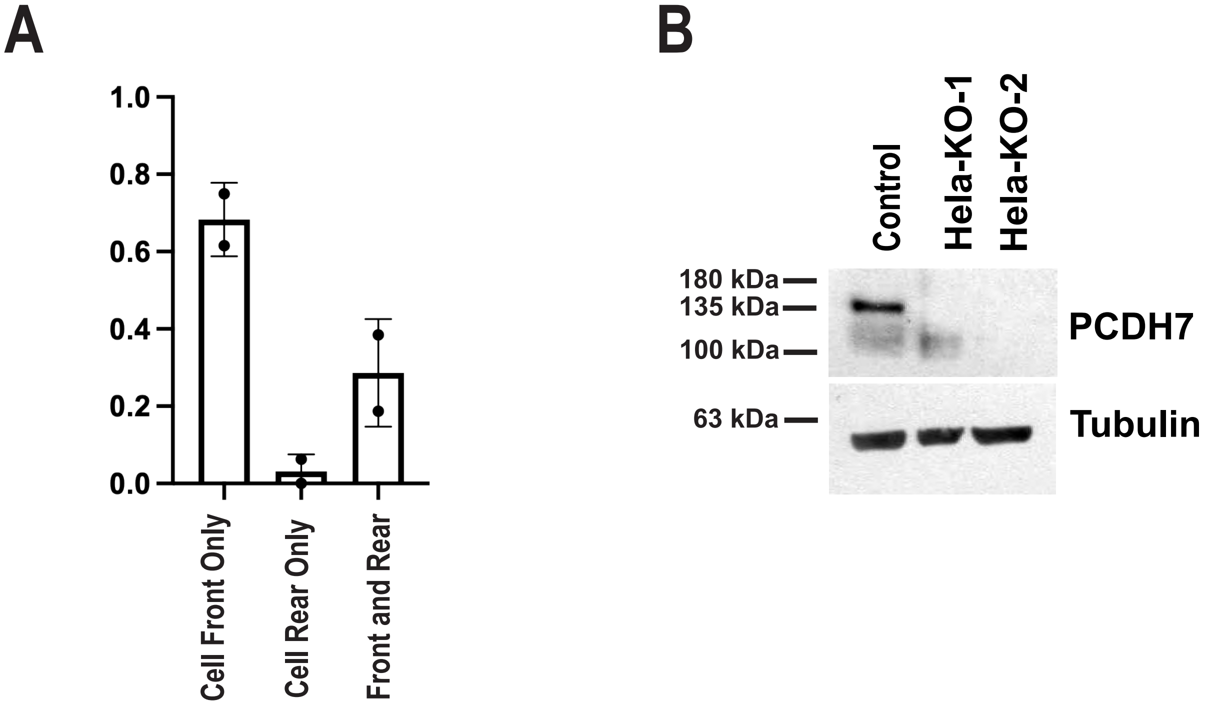

Qureshi et. al., Figure S2

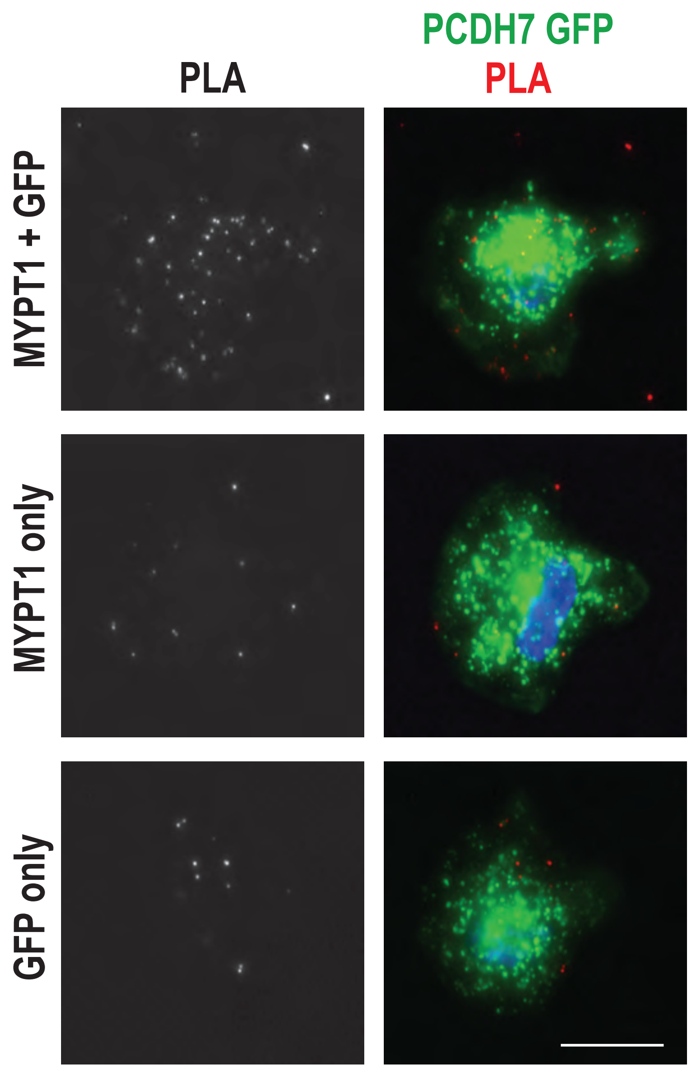

Qureshi et. al., Figure S3

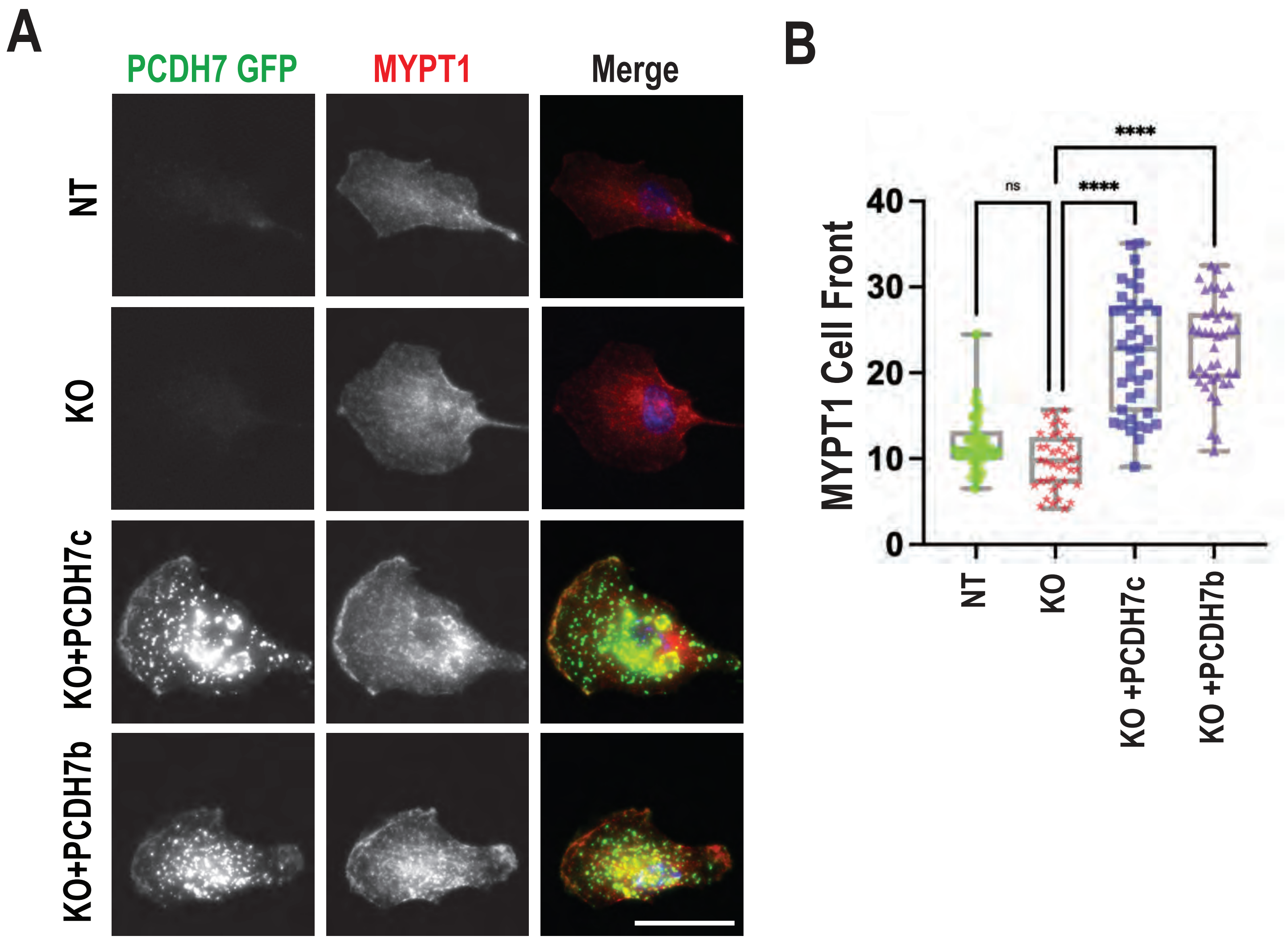

Qureshi et. al., Figure S4

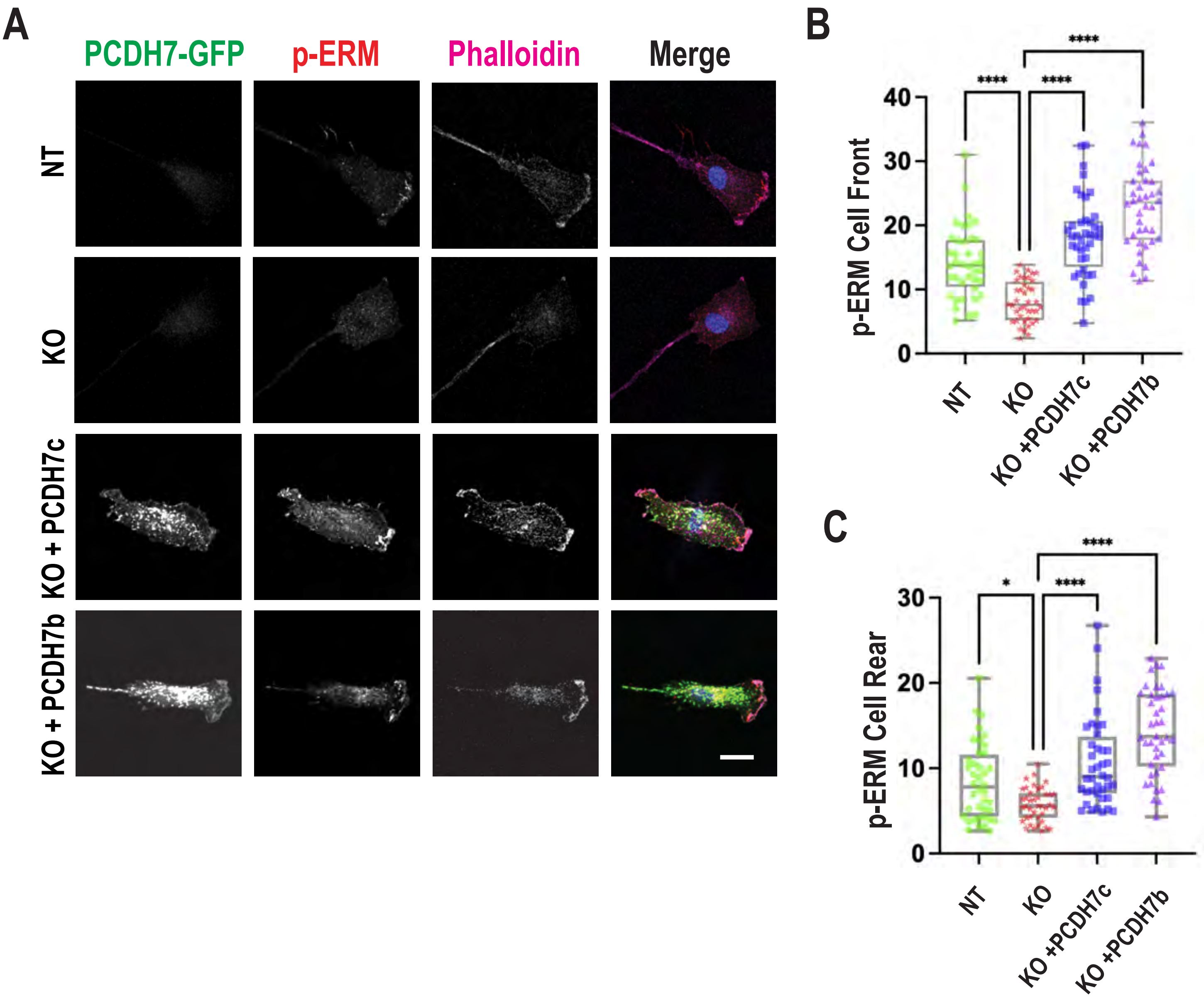

Qureshi et. al., Figure S5

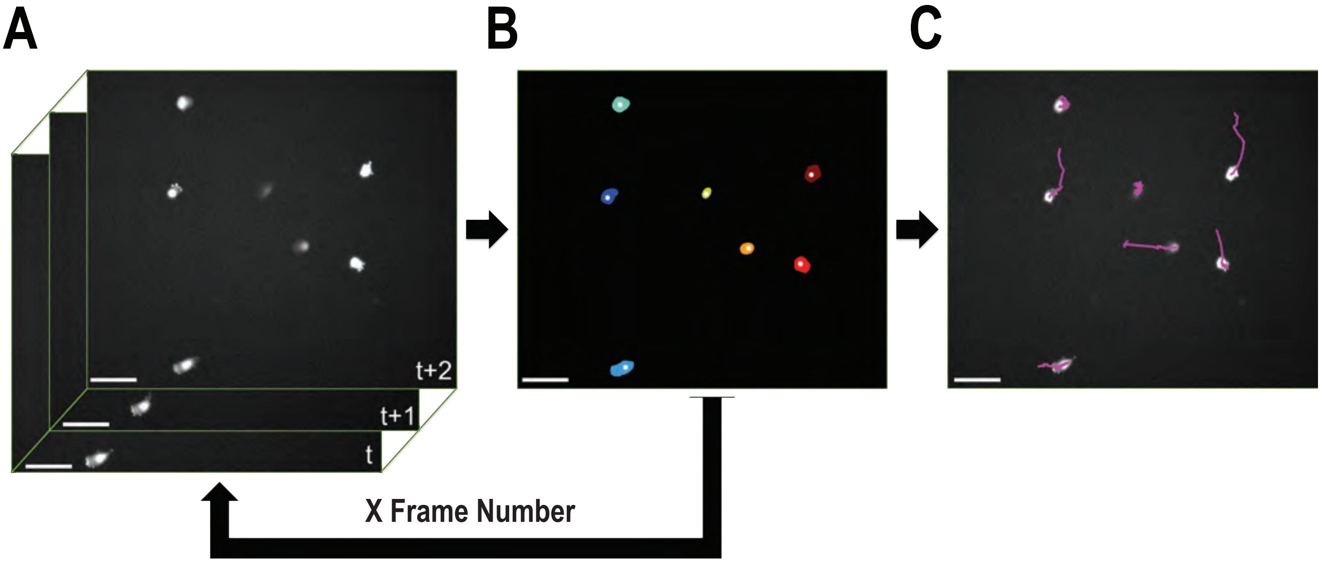
